# Identification of a key gain-of-function residue for effector binding by *in vitro* shuffling of barley *Mla NLR* genes

**DOI:** 10.1101/2024.10.27.619561

**Authors:** Xiaoxiao Zhang, Jialing Gao, Lucy M. Molloy, Lauren M. Crean, Simon J. Williams, John P. Rathjen

## Abstract

Natural plant populations maintain high resistance (*R*) gene diversities that provide effective pathogen resistance; however, such diversity has been reduced significantly by the genetic bottlenecks associated with plant domestication and breeding. Agricultural crops typically contain limited *R* gene diversity so resistance is often short-lived as pathogens evolve rapidly to evade recognition. The *mildew resistance locus A* (*Mla*) *R* gene family of barley and wheat represents a rich source of natural genetic variation that is ideal for mining disease resistance specificities. *Mla R* genes encode immune receptor proteins of the nucleotide-binding leucine-rich repeat (NLR) class that recognise unrelated plant pathogens by binding secreted virulence proteins termed effectors. The barley NLRs MLA13 and MLA7 confer resistance to different strains of the barley powdery mildew pathogen through direct interaction with the pathogen effectors AVR_A13_ and AVR_A7_ respectively. Using DNA shuffling, we generated a variant library by recombining the *Mla7* and *Mla13* genes *in vitro*. The variant library was cloned into yeast generating ∼4,000 independent clones and was screened for interaction with AVR_A13_ and AVR_A7_ using a yeast-two-hybrid (Y2H) assay. This yielded a number of clones that encode NLR proteins that interacted with AVR_A13_. Sequence analysis showed that the interacting MLA proteins can be clustered into three groups, all of which contain a critical residue from MLA13. While MLA13 and MLA7 differ by 30 residues across the LRR domain, the replacement of leucine to serine at this position in MLA7 facilitated interaction with AVR_A13_ in yeast and AVR_A13_-dependent immune signalling *in planta*. We have established a pipeline that evolves MLAs to recognise distinct pathogen effectors without the requirement for protein structural knowledge and the use of rationale design. We suggest these findings represent a step towards evolving novel recognition capabilities rapidly *in vitro*.

## Introduction

Plants possess a sophisticated immune system which relies on resistance (*R*) genes to detect and defend against pathogen invasions (Dodds & Rathjen, 2010). Natural *R* genes are numerous, diverse, and effective against a wide range of diseases. However, genetic bottlenecks associated with crop domestication and breeding have significantly reduced *R* gene diversity in crop species. Disease resistance conferred by a limited pool of *R* genes can be easily overcome by rapidly-evolving pathogens so is often short-lived. It is therefore essential to increase the *R* gene diversity available for breeding disease resistant crops.

A major class of *R* genes encodes nucleotide-binding leucine-rich repeat (NLR) proteins, which act intracellularly as receptors by recognising pathogen-secreted virulence proteins termed effectors (Dodds & Rathjen, 2010). Plant NLRs have characteristic structures comprising either a coiled-coil (CC) or Toll/interlukin-1 receptor/resistance (TIR) enzyme domain at the amino-terminus, a central nucleotide binding (NB) domain and a leucine-rich repeat (LRR) domain at their carboxy-termini. They can interact with pathogen effectors directly, or indirectly via accessory proteins. Direct interaction often occurs via the LRR domains or through non-canonical integrated domains (IDs) that exist in some NLRs. Effector recognition is typically highly specific. The LRR domains and IDs are therefore under positive selection pressure and demonstrate high sequence diversification compared to the CC, TIR or NB domains (Seeholzer *et al*., 2010; Bailey *et al*., 2018). Effector recognition typically leads to assembly of signalling complexes termed resistosomes (Huang *et al*., 2023) and immune activation is visible as localised cell death.

The *mildew resistance locus a* (*Mla*) gene family of cereals harbours great diversity, evident in the number of genes identified from various plant species and the diverse pathogen effectors that they recognise (Seeholzer *et al*., 2010). NLRs encoded by *Mla* genes possess a typical CC-NB-LRR structure. All MLA NLR proteins characterised to date interact with their corresponding effectors via direct interaction through their LRR domains (Chen *et al*., 2017; Saur *et al*., 2019; Bauer *et al*., 2021). There are 28 *Mla* genes at the barley *Mla* locus that confer resistance to various isolates of the fungal pathogen *Blumeria graminis* f. sp *hordei* (*Bgh*), the causative agent of powdery mildew disease (Seeholzer *et al*., 2010). The barley genes encoding the MLA1, MLA6, MLA7, MLA9, MLA10, MLA13 and MLA22 proteins share over 92% nucleotide (nt) sequence identity and recognise the *Bgh* effectors AVR_A1_, AVR_A6_, AVR_A7_, AVR_A9_, AVR_A10_, AVR_A13_ and AVR_A22_ respectively (Bauer *et al*., 2021). These effectors share an RNase-like protein fold despite <45% nt sequence identity. Genes within the barley *Mla* locus also confer resistance to the wheat stripe rust fungus *Puccinia striiformis* f. sp. *tritici* (Bettgenhaeuser *et al*., 2021) and the rice blast fungus *Magnaporthe oryzae* (Brabham et al., 2024). Recognition of *M. oryzae* is mediated by interaction between MLA3 and a MAX-fold effector (de Guillen *et al*., 2015). The wheat *Mla* genes *Sr50* and *Sr33* confer resistance to stem rust disease caused by the fungus *Puccinia graminis* f. sp. *tritici* (*Pgt*) (Periyannan *et al*., 2013; Mago *et al*., 2015). The *Pgt* effector AvrSr50 that is recognised by Sr50 shares structural similarity with cupin superfamily members and carbohydrate hydrolases (Ortiz *et al*., 2022). A large number of *Mla* genes were also found in the wild barley species *Hordeum spontaneum* (Maekawa *et al*., 2019), and an *MlaR* gene called *TmMla1* was identified from the diploid wheat species *Triticum monococcum* (Jordan *et al*., 2011). More recently, the rice *Mla* gene *RYMV3* was found to confer resistance to rice yellow mottle virus through recognition of the viral coat protein (Bonnamy *et al*., 2023). *RYMV3* shares only ∼54% nt sequence identity with barley *Mla* genes (compared to 85% between barley and wheat *Mlas*) so this new discovery significantly expands the potential recognition spectrum of MLA NLRs.

The broad recognition spectrum mediated by the *Mla* family offers great opportunities for mining and engineering disease resistance specificities. NLR proteins have been engineered through rational design and targeted mutagenesis, exploiting natural amino acid polymorphisms informed by protein structures (Tamborski *et al*., 2023; Lawson *et al*., 2024). Directed evolution methods can be used to rapidly increase *NLR* gene diversity (Greenwood *et al*., 2023). These include *in vitro* gene recombination to mimic the natural evolution of modular proteins, which is especially suitable for recombining natural diversity. However, despite pilot studies with the potato *Rx* and *R3a NLR* genes (Harris *et al*., 2013; Chapman *et al*., 2014), an efficient pipeline to exploit the natural diversity of *NLR* genes has not yet been established.

Here we report the *in vitro* recombination of closely-related barley *Mla* genes and more distantly related wheat *Mla* genes using DNA shuffling. The recombinant proteins were screened for direct interaction with *Bgh* effectors using yeast-two-hybrid assays and tested for the cell death immune response *in planta*. Our study identified three groups of *Mla13* and *Mla7* recombinants and demonstrated their ability to recognise *Bgh* AVR_A13_ variants. These recombinants led to the identification of a gain of function mutantion in *Mla7*. Our work provides a path to evolving *Mla* genes to expand their ability to recognise pathogens.

## Results and discussion

### *In vitro* recombination of *Mla* genes via DNA shuffling

To explore *in vitro* gene recombination among different *Mla* genes, DNA shuffling was conducted pairwise among barley *Mla7, Mla13, Mla10* and *Mla1*, and wheat *Sr33, TmMla1* and *Sr50*. The corresponding pathogen *AVR* genes have been identified for all of the selected barley *Mla* genes (Lu *et al*., 2016; Saur *et al*., 2019) and for wheat *Sr50* (Chen et al., 2017), enabling interaction studies. DNA shuffling follows a three-step protocol from gene fragmentation through recombination of fragments and then amplification of the final products (Stemmer, 1994). The efficiency of *in vitro* recombination during DNA shuffling is indicated by the final yield. Self-shuffling of one gene generated a single PCR product of the correct size, whereas shuffling of two genes with <80% nt sequence identity generated multiple PCR products and showed lower target to non-specific product ratios (Stemmer, 1994; Abecassis *et al*., 2000). DNA shuffling of the *Mla* genes generated multiple PCR products with various target to non-specific product ratios (Figure 1A and Figure S1). High yields were obtained from the self-shuffling of *Mla13* and the pairwise shuffling of barley *Mla* genes that share 95-98% nt sequence identities, as indicated by a dominant PCR product of the expected ∼3,000 base pair (bp) size. A further shuffling of *Mla1, Mla7, Mla9, Mla10, Mla13 and Mla22* generated a high yield similar to the pairwise shuffling between barley *Mla* genes (Figure S1). Pairwise shuffling between wheat *Mla* genes that share 86-90% nt sequence identities led to intermediate yields, as indicated by a 3,000 bp PCR product with slightly higher intensity than the non-specific bands. Shuffling of barley and wheat *Mla* genes generated intermediate to low yields, where the yield of the target product was equal to or less than the non-specific products, despite all sharing approximately 85% sequence identities. For example, when shuffled with *Mla13, Sr33* and *TmMla1* showed intermediate product yields, while Sr50 showed low product yield. Phylogenetic analysis revealed that *Sr33* and *TmMla1* cluster in the same subgroup, separate from *Sr50* (Figure 1B). These results suggest that efficiency of DNA shuffling between *Mla* genes depends on the gene sequence identities and their phylogenetic relationships.

**Figure 1.**
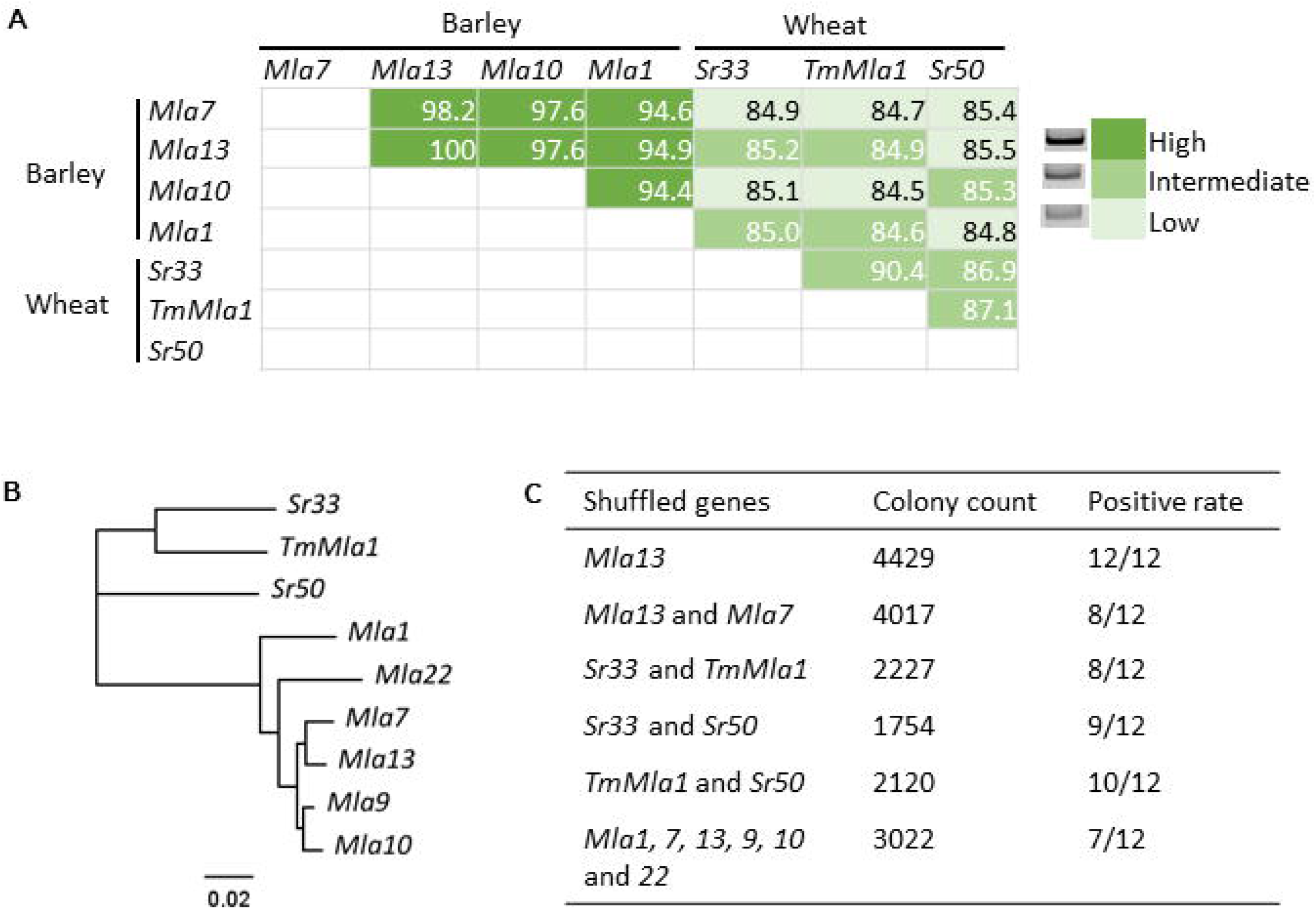
Pairwise DNA shuffling between barley and wheat *Mla* genes. (A) Sequence alignment matrix for selected *Mla* genes. The numbers indicate the percentage nucleotide sequence identity between each pair. The genes were shuffled and amplified in a pairwise fashion and the DNA products analysed by gel electrophoresis. The yields of each amplification were assessed by gel band intensity. Representative bands are shown on the right and approximate yields are indicated by green shades. (B) Neighbor-Joining phylogenetic tree of the nine selected *Mla* genes based on nucleotide sequence alignment. The scale bar shows estimated substitutions per site (C) Yeast expression libraries generated from the DNA shuffling experiments. Colony count indicates the number of colonies generated by transformation of the recombinant genes into yeast. Positive rate indicates the number of full-length genes identified by colony PCR for 12 colonies selected from each library.

To create expression libraries containing the recombinant genes, the shuffled DNA products were cloned into the Y2H vector pGBDT7 using *in vivo* recombinational cloning in yeast. The resulting libraries express MLA variants fused to an N-terminal GAL4 BD domain (BD-MLA). Six yeast libraries were created (Figure 1C). Each library generated ∼4,000 yeast colonies from shuffling between barley *Mla* genes and ∼2,000 colonies from shuffling between wheat *Mla* genes, which is consistent with the DNA yields of the shuffling experiments. Colony PCR indicated successful transformation of the recombinant genes into yeast, for which over half of the selected colonies contained a full-length *Mla* gene, except for *Mla13* self-shuffling in which all of the selected colonies contained the *Mla13* gene (Figure S2). This demonstrates successful shuffling of full-length *Mla NLR* genes with 85-98% nt sequence identities. The method can easily scale to produce larger numbers of recombinant clones in yeast.

### Recombinant *Mla* genes encode variants that interact with AVR_A13_

The barley MLA7 and MLA13 proteins interact directly with two AVR_A7_ (AVR_A7_-1 and AVR_A7_-2) and four AVR_A13_ (AVR_A13_-V2, AVR_A13_-1, AVR_A13_-2 _and_ AVR_A13_-3) effector variants respectively, as shown by Y2H assays (Lu *et al*., 2016; Saur *et al*., 2019). The shuffled *Mla13* + *Mla7* library termed *Mla13/7* was screened for interactions with AVR_A13_-V2 or AVR_A7_-2 fused to an N-terminal GAL4 AD domain (AD-AVR) using a modified mating-based method. Using this Y2H system, interactions between MLA13 and AVR_A13_-V2 and Sr50 and AvrSr50 were detected, but we were not able to detect interaction between MLA7 and AVR_A7_-2 (Figure 2A), despite confirmation of protein accumulation for both MLA7 and AVR_A7_-2 (Figure S3). MLA13 did not show interaction with negative controls including the AD domain empty vector (EV) and AD domain fused to the unrelated TIR domain of the flax NLR protein L6 (L6TIR) (Bernoux *et al*., 2011). Previous studies using a LexA Y2H system identified an interaction between MLA7 and AVR_A7_-2 that was noticeably weaker than that between MLA13 and AVR_A13_-V2 (Saur *et al*., 2019). The GAL4 Y2H system may not be sensitive enough to detect the MLA7 and AVR_A7_-2 interaction. We therefore focused on screening the *Mla13/7* library for AVR_A13_-V2 interactors.

**Figure 2.**
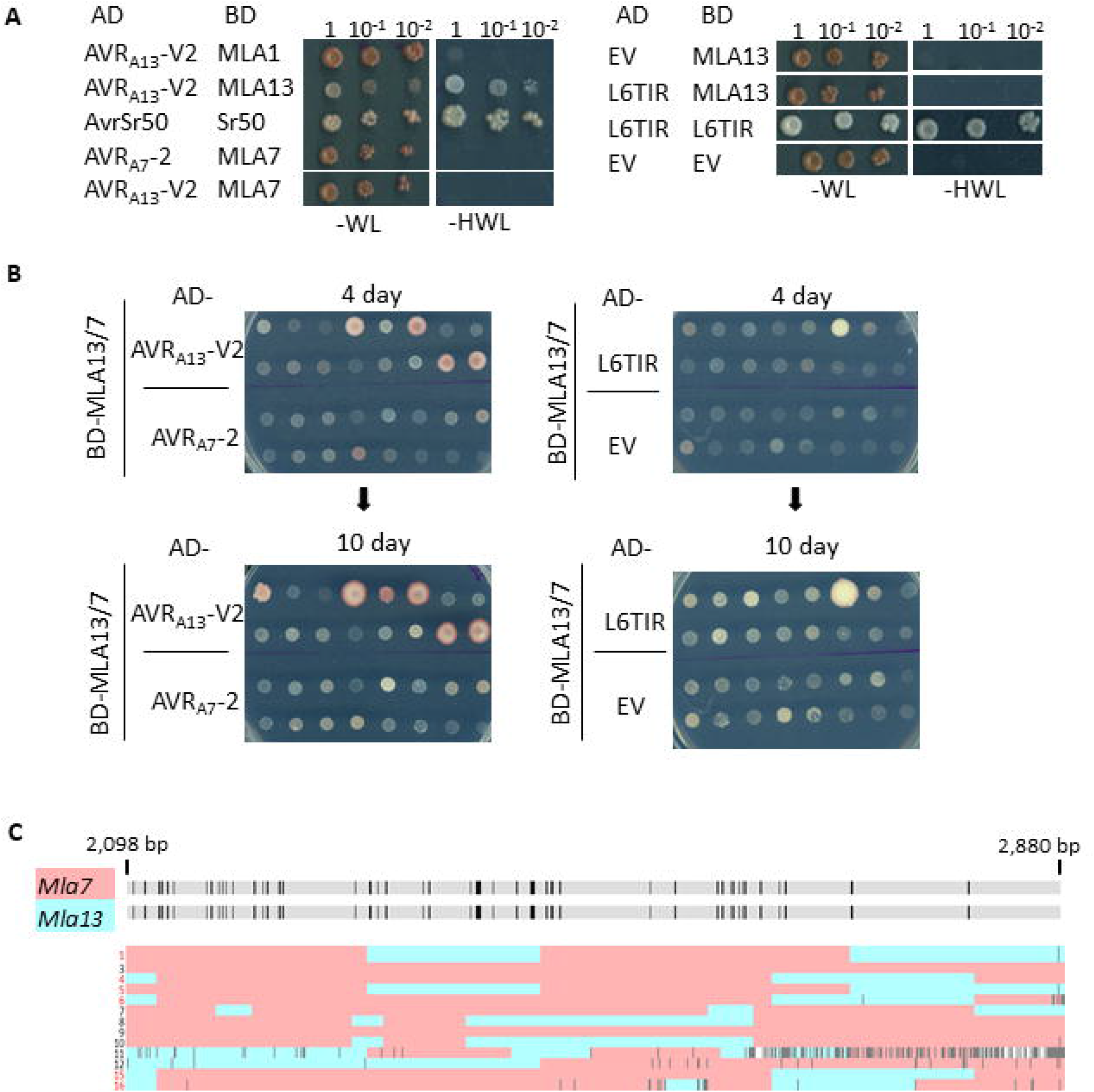
Yeast-two-hybrid screening of the recombinant gene libraries for interaction with AVR proteins. (A) Positive controls for MLA-AVR Y2H interactions. Diploid yeast strains carrying both the GAL4 activation domain (AD) and the GAL4 binding domain (BD) constructs were grown on synthetic drop-out (SD) media lacking tryptophan and leucine (-WL), and interactions between proteins were detected by growth on SD media additionally lacking histidine (-HWL). EV denotes empty vector. (B) Growth of 16 selected colonies from each mating of the MLA13/7 library (BD-MLA13/7) against AVR proteins and controls on SD-HWL. Top panel: growth at 4 days; bottom panel: growth at 10 days. (C) Sequence analysis of shuffled *Mla13/7* sequences recovered from diploid colonies mated with AD-AVR_A13_-V2. The upper panel shows sequence alignment of *Mla7* and *Mla13* in the 3’-terminal 2,098-2,880 bp region encoding the LRR domains. Variable sites are highlighted in dark grey and conserved sites are in light grey. The lower panel is a cartoon depiction of sequence alignment and DNA shuffling analysis of 13 recombinant sequences. The colony number that the sequence was derived from is shown on the left, with AVR_A13_-V2 interacting colonies highlighted in red. The gene fragments assigned to *Mla7* and *Mla13* are coloured in red and blue respectively. Mutations that were not assigned to either parent are shown in dark grey. White regions denote where sequences could not be assigned to either parent due to mutations at the variable sites.

For library screening, a small-scale screen was first carried out in which the BD-MLA13/7 yeast library was mated separately with strains containing the effector constructs AD-AVR_A13_-V2 or AD-AVR_A7_-2, or the negative controls AD-L6TIR or AD-EV. Diploid yeast colonies carrying both the BD- and AD-constructs were randomly picked from each combination and tested for interaction as indicated by growth on selective plates over 4 to 10 days. Significant yeast growth was found for six out of 16 sampled colonies (6/16) mated with AD-AVR_A13_-V2, and for one colony (1/16) mated with AD-L6TIR, but none in the AD-AVR_A7_-2 or AD-EV matings (0/16) (Figure 2B). Colony PCR showed that most of the selected colonies contained an insert of 3,000 bp, equivalent to a full-length *Mla* gene (Figure S4A). However, the single positive colony from the AD-L6TIR interaction plate contained an insert of 600 bp, which may explain activation of the reporter gene. The recombinant *Mla13/7* genes were then amplified from the 16 colonies mated with AD-AVR_A13_-V2 and the LRR encoding regions were sequenced using Sanger sequencing, after which 13 sequences (Figure S4B) were recovered and assembled using *Mla13* as a reference. *Mla13* and *Mla7* are conserved within the first 2,000 bp and vary in the 3’ 880 bp encoding the LRR domain. Sequence analysis of the variable region showed that two sequences represent the WT *Mla7* gene and the remaining 11 sequences contain fragments from both *Mla7* and *Mla13* correlating to 1-3 gene recombination events per gene within the region (Figure 2C).

A larger scale screen was next carried out in which diploid (mated) yeast colonies were replica plated onto the interaction plates. This resulted in growth of ∼280 colonies for AD-AVR_A13_-V2 and ∼40 colonies for AD-AVR_A7_-2. Colony PCR identified 16 full-length *Mla* genes out of 40 colonies picked from the AD-AVR_A13_-V2 interaction plate. Only one of 40 colonies from the AD-AVR_A7_-2 interaction plate contained a full-length *Mla* gene, however it was not able to grow on the replica interaction plate, indicating a false positive (Figure S4C). This indicates that MLA13/7 variants capable of interacting with AVR_A13_-V2 were identified, but none for AVR_A7_-2.

Sequence alignment of the 21 positive *Mla13/7* clones generated from the two batches of screening re-ordered the recombinant genes into 9 clusters and showed reduced sequence diversity compared to the non-selected library (Figure 2C and S5A). Sequences from each screen segregated in the sequence alignment which may be due to difference in the quality of Sanger sequencing. All of the positive genes from Batch 1 were selected except for cluster 4 that contains a synonymous variation, and representative genes were selected from each cluster of Batch 2. The corresponding plasmids were recovered and full-length sequences obtained using Nanopore whole plasmid sequencing. Sequences were recovered for 9 of the 10 selected recombinant genes (R1-R10; sequencing failed for R2). Sequence alignments showed that the recombinant genes clustered into three sub-groups (Figure 3A). Group A contains two clones (R1 and R3) representing the *Mla7* sequence with the regions between bases 2,299-2,452 and the 3’-end bases 2,704-2,880 exchanged with the reciprocal *Mla13* sequences. This corresponds to 11 amino acid (aa) variations in the LRR domain between residues 767-812 and at residues 902 and 935. Strikingly, both clones contain a point mutation at base 892, substituting residue 298 from arginine (R) in both MLA13 and MLA7 to glycine (G). The R298 residue is located in the NB domain conserved in all barley MLAs and the corresponding positions in wheat Sr50 and Sr33, but the cognate residue is a serine in TmMLA1 (Figure S5B). A survey using NCBI BLAST identified a methionine at the corresponding position in three wild barley *Mla* gene candidates and an asparagine in one *Mla* candidate gene from the wild wheat *Dasypyrum villosum*. Our DNA shuffling experiments apparently introduced an untemplated mutation at this position which was selected in the Y2H screen. Further investigation is required to determine whether the R298G mutation co-varies with the additional amino acid variations between residues 767-812 in the LRR domain. Group B containing five clones (R4, R5, R6, R8 and R10) represents the *Mla7* sequence with the bases 2,100-2,122 and 2,645-2,803 replaced with the reciprocal sequences from *Mla13*. This corresponds to four aa variations, at residues 700, 704, 880 and 902. Group C containing two clones (R7 and R11) represents the *Mla7* sequence with the regions between bases 2,100-2,162 and 3’-end bases 2,704-2880 replaced by reciprocal sequences from *Mla13*. This corresponds to seven aa variations between residues 700-712, and at residues 902 and 935. All of the recombinant genes retrieved during the screen contain the 2,704-2,802 bp region from *Mla13* which corresponds to a single variable aa residue at position 902. *Mla13* and the selected genes encode a serine (S) residue at this position, while *Mla7* encodes a leucine (L) residue, suggesting an important role for S902 in the MLA interaction with AVR_A13_-V2.

**Figure 3.**
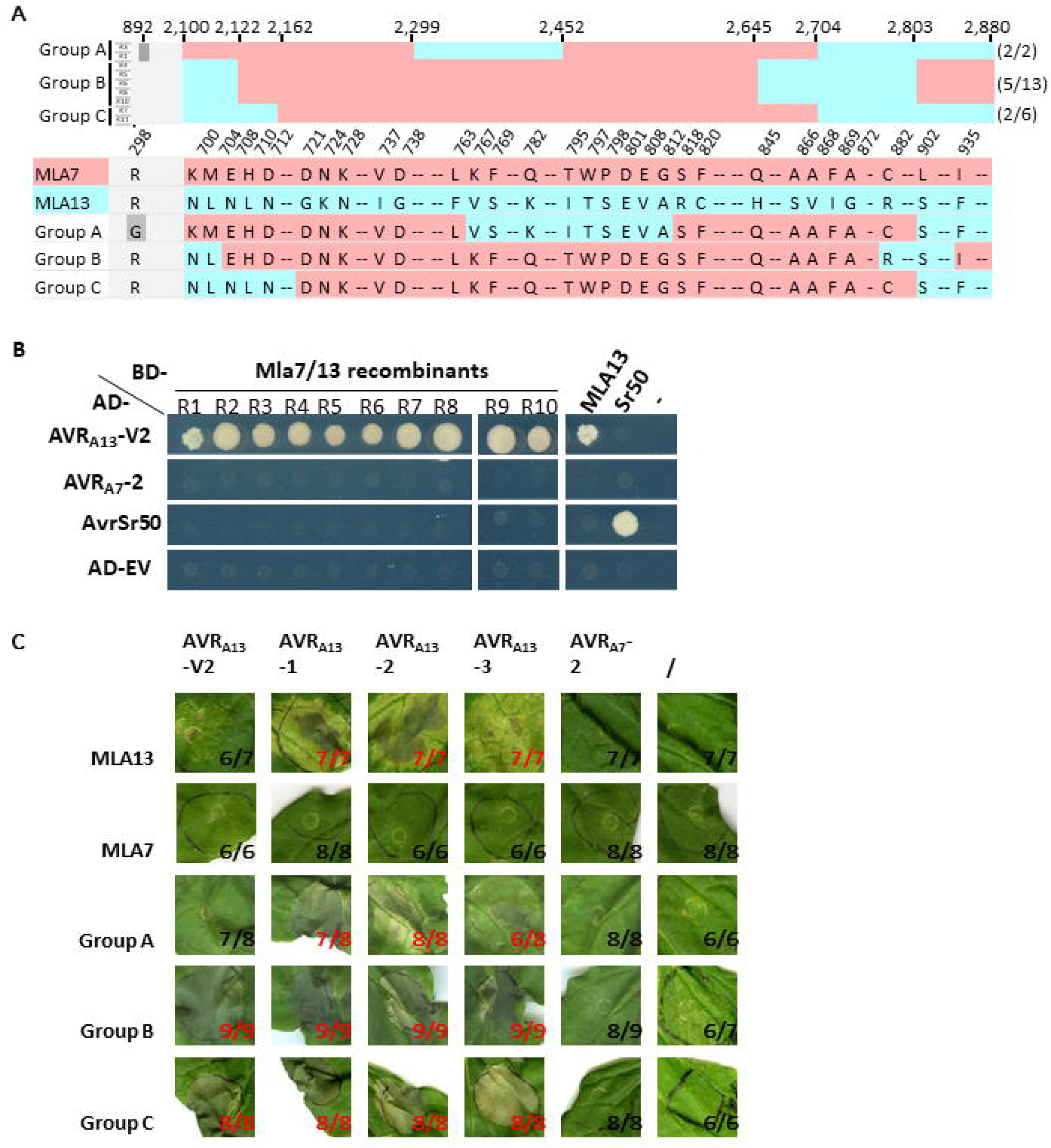
AVR interaction and cell death induction by recombinant *Mla13/7* genes. (A) Sequence analysis of recombinant *Mla13/7* genes encoding variants that interacted with AVR_A13_-V2 in yeast. The upper panel shows a cartoon depicting nucleotide sequence alignments and the lower panel shows the variant amino acids. The sequences assigned to *Mla7* and *Mla13* are coloured in red and blue respectively. Conserved regions are shown in light grey and the mutation that is not found in either parent is highlighted in dark grey. The selected *Mla13/7* genes were clustered into three groups A-C based on sequence characteristics. The numbers (x/y) indicate the numbers of clones sequenced with Nanopore sequencing of the total number of clones sequenced with Sanger sequencing for each group. The conserved amino acid sequences are either not represented or shown as dashes. (B) Y2H assays for recombinant MLA13/7 variants against AVR_A13_-V2, AVR_A7_-2 and AvrSr50. Yeast cells were co-transformed with *Mla13/7* recombinant genes, *Mla13* or *Sr50* fused to *GAL4 BD* or water control (-), and *AVR* genes fused to *GAL4 AD*. Protein interactions were detected via yeast growth on SD-HWL media. (C) Cell death assays for MLA13/7 variants in *Nicotiana benthamiana* leaves. Agrobacterium strains carrying MLA13, MLA7 or one of the MLA13/7 variants were co-infiltrated with Agrobacterium strains carrying one of the AVR_A13_ variants, AVR_A7_-2 or alone (/) into *N. benthamiana* leaves. Images were taken 5 days after infiltration. Yellow to dark spots on the leaves indicate cell death. The numbers indicate the numbers of leaves showing the representative phenotypes of the total number of leaves infiltrated. The numbers on leaves showing cell death phenotypes are coloured in red.

### MLA13/7 variants interact with AVR_A13_-V2 in yeast and trigger AVR_A13_-dependent cell death in *Nicotiana benthamiana*

Independent Y2H tests to validate interactions between the MLA13/7 variants and AVR_A13_-V2 were conducted by co-transformation of the respective GAL4 BD and AD constructs into yeast. Sr50 and AvrSr50 were used as controls because we were not able to detect MLA7 and AVR_A7_-2 interaction using the GAL4 Y2H system. The ten selected MLA13/7 variants showed interaction with AVR_A13_-V2, but not AVR_A7_-2, AvrSr50 or AD-EV (Figure 3B). Interaction was detected for the positive controls MLA13 and AVR_A13_-V2, and Sr50 and AvrSr50. Therefore, the results from the initial screen were confirmed in the independent tests.

MLA13 triggers localised cell death when expressed transiently in *N. benthamiana* in combination with AVR_A13_-1, AVR_A13_-2 and AVR_A13_-3, but not with AVR_A13_-V2 (Saur *et al*., 2019) (Figure 3C and Figure S6). Similarly, we found that Group A MLA13/7 variants triggered cell death when co-expressed with AVR_A13_-1, AVR_A13_-2 and AVR_A13_-3, but not with AVR_A13_-V2 (Figure 3C and Figure S6). Despite this, Group B and Group C MLA13/7 variants triggered cell death with all four AVR_A13_ variants. None of the MLA13/7 variants, MLA13 or MLA7 triggered cell death with AVR_A7_-2 or alone, despite confirmation of similar protein accumulation for MLA13, MLA7 and all AVR_A_s (Figure S7). Therefore, Group A variants retained the recognition specificities of MLA13, while Group B and C variants gained recognition of AVR_A13_-V2 in *N. benthamiana*.

### A single mutation enables MLA7 to interact with Avr_A13_-V2 in yeast and trigger cell death *in N. benthamiana*

To further investigate the role of Ser-902 in AVR recognition, an MLA7 mutant was generated to substitute L902 with serine, designated MLA7^L902S^. MLA7^L902S^ but not MLA7 interacted with AVR_A13_-V2 in yeast (Figure 4A) and triggered cell death when co-expressed with AVR_A13_-V2, AVR_A13_-1 or AVR_A13_-2 in *N. benthamiana* (Figure 4B and Figure S8). Natural sequence polymorphism at this position has been found to alter *Mla13* function (Bettgenhaeuser *et al*., 2021). A recent study revealed that S902 is located at the periphery of the interaction interface between MLA13 and AVR_A13_-1, suggesting that it may not participate directly in the interaction (Lawson *et al*., 2024). However, MLA7^L902S^ gained recognition of AVR_A13_-1 and AVR_A13_-V2, and retained recognition of AVR_A7_-1 and AVR_A7_-2 in cell death assays (Lawson *et al*., 2024). Thus our findings independently support the conclusions of the structural study.

**Figure 4.**
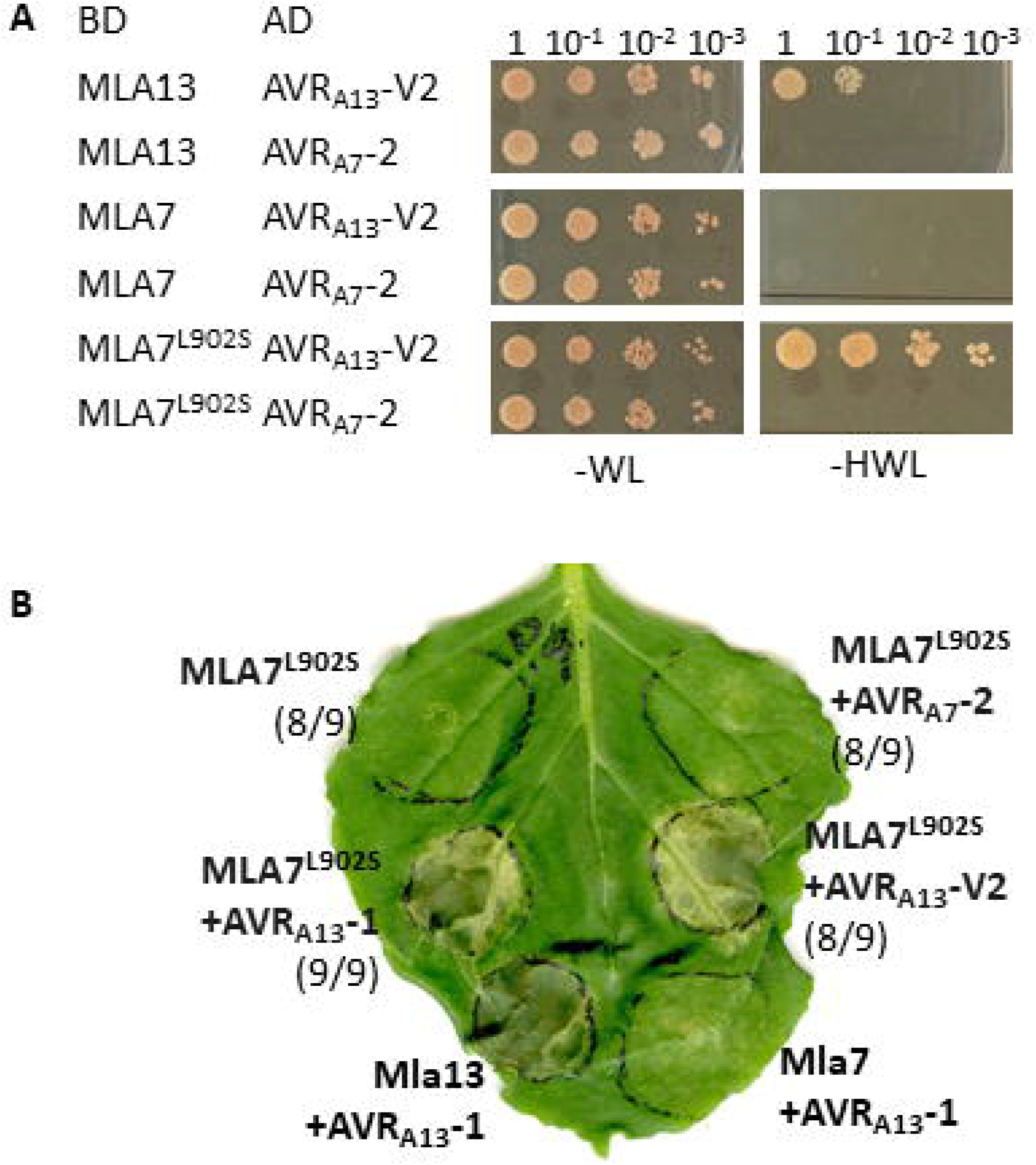
Interaction and cell death studies of the *Mla7*^*L902S*^ mutant. (A) Yeast interaction assays for the *MLA7*^*L902S*^ mutant. Yeast cells were co-transformed with *Mla7*^*L902S*^ fused to *GAL4 BD* and *AVR* genes fused to *GAL4 AD*. Diploid yeast strains carrying both the *AD-* and *BD-* constructs were selected based on yeast growth on SD-WL media and protein interactions were detected by growth on SD-HWL media. (B) Cell death assays for *Mla7*^*L902S*^ in *N. benthamiana*. Agrobacterium strains carrying *Mla13, Mla7* or *Mla7*^*L902S*^ were co-infiltrated with Agrobacterium strains carrying *AVR*_*A13*_*-V2, AVR*_*A13*_*-1* or *AVR*_*A7*_*-2* into *N. benthamiana* leaves. Images were taken five days after infiltration. Cell death is indicated by yellow to dark spots on the leaves. The numbers indicate the numbers of leaves showing the representative phenotypes of the total number of leaves infiltrated.

## Conclusions

NLR engineering is an emerging avenue for manipulating host recognition specificities of distinct pathogen AVRs, which holds promise for building novel disease resistance traits in crop plants. Schemes that utilise rational design based on real or modelled effector-NLR structures represent a popular strategy (Greenwood *et al*., 2023). Here we show how a non-targeted approach exploiting natural sequence diversity can be applied to NLR engineering. Our study demonstrates the successful application of *in vitro* DNA shuffling to rapid generation and identification of functional NLR variants from the natural *Mla* family. We identified an MLA variant with altered AVR recognition capabilities which not only supports the observations of a recent structural study (Lawson *et al*., 2024) but was generated independently of structural knowledge or prior consideration of amino acid polymorphisms. This is an important step towards evolving NLRs to detect unrecognised pathogen effectors, which is not possible through rational design. The diversity of the variant library is currently limited because the parents we used are closely-related barley genes. Screening of this library thus only altered the recognition of known AVRs. We have successfully shuffled more distantly-related *Mla* genes from wheat and barley, which will generate variant libraries with higher diversity that may allow expansion of the spectrum of effector recognition. In addition, we established a Y2H method that enables high throughput screening against a pool of effector proteins. Together, we believe this represents a significant step to evolving NLR recognition specificity *in vitro*.

## Author contributions

XZ and JPR planned and designed the research study. XZ, JG, LMM and LMC performed the experiments; XZ, SJW and JPR analysed the data. XZ wrote the original draft and all authors contributed to writing, reviewing and editing of the manuscript.

## Data and materials availability

Where possible materials generated in this study will be made available upon request to the corresponding authors. All other data are available in the main text or supplementary materials.

## Acknowledgments

We thank Isabel Saur for her insightful discussion and suggestion. The entry constructs of *Mla13*/*AVR*_*A13*_ and *Mla7*/*AVR*_*A7*_ were kindly provided by the Schulze-Lefert group (MPIPZ, Germany). We thank the Plant Services Team at the Australian National University for providing *N. benthamiana* seedlings. This work was funded by the Australian Research Council (ARC) Discovery Early Career Researcher Award (DE210100323) and the Australian Academy of Science J G Russell Award both to XZ. SJW was a recipient of the ARC Future Fellowship (FT200100135). Work in JPR’s laboratory is supported by the ARC Discovery Project grant (DP190103040).

## References

Abecassis V, Pompon D, Truan G. 2000. High efficiency family shuffling based on multi-step PCR and in vivo DNA recombination in yeast: statistical and functional analysis of a combinatorial library between human cytochrome P450 1A1 and 1A2. Nucleic Acids Res 28(20): E88.

Bailey PC, Schudoma C, Jackson W, Baggs E, Dagdas G, Haerty W, Moscou M, Krasileva KV. 2018. Dominant integration locus drives continuous diversification of plant immune receptors with exogenous domain fusions. Genome Biol 19(1): 23.

Bauer S, Yu D, Lawson AW, Saur IML, Frantzeskakis L, Kracher B, Logemann E, Chai J, Maekawa T, Schulze-Lefert P. 2021. The leucine-rich repeats in allelic barley MLA immune receptors define specificity towards sequence-unrelated powdery mildew avirulence effectors with a predicted common RNase-like fold. PLoS Pathog 17(2): e1009223.

Bernoux M, Ve T, Williams S, Warren C, Hatters D, Valkov E, Zhang X, Ellis JG, Kobe B, Dodds PN. 2011. Structural and functional analysis of a plant resistance protein TIR domain reveals interfaces for self-association, signaling, and autoregulation. Cell Host Microbe 9(3): 200–211.

Bettgenhaeuser J, Hernandez-Pinzon I, Dawson AM, Gardiner M, Green P, Taylor J, Smoker M, Ferguson JN, Emmrich P, Hubbard A, Bayles R, Waugh R, Steffenson BJ, Wulff BBH, Dreiseitl A, Ward ER, Moscou MJ. 2021. The barley immune receptor Mla recognizes multiple pathogens and contributes to host range dynamics. Nat Commun 12(1): 6915.

Bonnamy M, Pinel-Galzi A, Gorgues L, Chalvon V, Hebrard E, Cheron S, Nguyen TH, Poulicard N, Sabot F, Pidon H, Champion A, Cesari S, Kroj T, Albar L. 2023. Rapid evolution of an RNA virus to escape recognition by a rice nucleotide-binding and leucine-rich repeat domain immune receptor. New Phytol 237(3): 900–913.

Brabham HJ, Gomez De La Cruz D, Were V, Shimizu M, Saitoh H, Hernandez-Pinzon I, Green P, Lorang J, Fujisaki K, Sato K, Molnar I, Simkova H, Dolezel J, Russell J, Taylor J, Smoker M, Gupta YK, Wolpert T, Talbot NJ, Terauchi R, Moscou MJ. 2024. Barley MLA3 recognizes the host-specificity effector Pwl2 from Magnaporthe oryzae. Plant Cell 36(2): 447–470.

Chapman S, Stevens LJ, Boevink PC, Engelhardt S, Alexander CJ, Harrower B, Champouret N, McGeachy K, Van Weymers PS, Chen X, Birch PR, Hein I. 2014. Detection of the virulent form of AVR3a from Phytophthora infestans following artificial evolution of potato resistance gene R3a. PLoS One 9(10): e110158.

Chen J, Upadhyaya NM, Ortiz D, Sperschneider J, Li F, Bouton C, Breen S, Dong C, Xu B, Zhang X, Mago R, Newell K, Xia X, Bernoux M, Taylor JM, Steffenson B, Jin Y, Zhang P, Kanyuka K, Figueroa M, Ellis JG, Park RF, Dodds PN. 2017. Loss of AvrSr50 by somatic exchange in stem rust leads to virulence for Sr50 resistance in wheat. Science 358(6370): 1607–1610.

de Guillen K, Ortiz-Vallejo D, Gracy J, Fournier E, Kroj T, Padilla A. 2015. Structure analysis uncovers a highly diverse but structurally conserved effector family in phytopathogenic fungi. PLoS Pathog 11(10): e1005228.

Dodds PN, Rathjen JP. 2010. Plant immunity: towards an integrated view of plant-pathogen interactions. Nat Rev Genet 11(8): 539–548.

Engler C, Youles M, Gruetzner R, Ehnert TM, Werner S, Jones JDG, Patron NJ, Marillonnet S. 2014. A Golden Gate modular cloning toolbox for plants. ACS Synth Biol 3(11): 839–843.

Greenwood JR, Zhang X, Rathjen JP. 2023. Precision genome editing of crops for improved disease resistance. Curr Biol 33(11): R650–R657.

Harris CJ, Slootweg EJ, Goverse A, Baulcombe DC. 2013. Stepwise artificial evolution of a plant disease resistance gene. Proc Natl Acad Sci U S A 110(52): 21189–21194.

Huang S, Jia A, Ma S, Sun Y, Chang X, Han Z, Chai J. 2023. NLR signaling in plants: from resistosomes to second messengers. Trends Biochem Sci 48(9): 776–787.

Jordan T, Seeholzer S, Schwizer S, Toller A, Somssich IE, Keller B. 2011. The wheat Mla homologue TmMla1 exhibits an evolutionarily conserved function against powdery mildew in both wheat and barley. Plant J 65(4): 610–621.

Kaiser C, Michaelis S, Mitchell A, Cold Spring Harbor Laboratory. 1994. Methods in yeast genetics : a Cold Spring Harbor Laboratory course manual. Cold Spring Harbor, NY: Cold Spring Harbor Laboratory Press.

Lawson AW, Flores-Ibarra A, Cao Y, An C, Neumann U, Gunkel M, Saur IML, Chai J, Behrmann E, Schulze-Lefert P. 2024. The barley MLA13-AVRA13 heterodimer reveals principles for immunoreceptor recognition of RNase-like powdery mildew effectors. bioRxiv: 2024.2007.2014.603419.

Lu X, Kracher B, Saur IM, Bauer S, Ellwood SR, Wise R, Yaeno T, Maekawa T, Schulze-Lefert P. 2016. Allelic barley MLA immune receptors recognize sequence-unrelated avirulence effectors of the powdery mildew pathogen. Proc Natl Acad Sci U S A 113(42): E6486–E6495.

Maekawa T, Kracher B, Saur IML, Yoshikawa-Maekawa M, Kellner R, Pankin A, von Korff M, Schulze-Lefert P. 2019. Subfamily-specific specialization of RGH1/MLA immune receptors in wild barley. Mol Plant Microbe Interact 32(1): 107–119.

Mago R, Zhang P, Vautrin S, Simkova H, Bansal U, Luo MC, Rouse M, Karaoglu H, Periyannan S, Kolmer J, Jin Y, Ayliffe MA, Bariana H, Park RF, McIntosh R, Dolezel J, Berges H, Spielmeyer W, Lagudah ES, Ellis JG, Dodds PN. 2015. The wheat Sr50 gene reveals rich diversity at a cereal disease resistance locus. Nat Plants 1: 15186.

Meyer AJ, Ellefson JW, Ellington AD. 2014. Library generation by gene shuffling. Curr Protoc Mol Biol 105: Unit 15 12.

Ortiz D, Chen J, Outram MA, Saur IML, Upadhyaya NM, Mago R, Ericsson DJ, Cesari S, Chen C, Williams SJ, Dodds PN. 2022. The stem rust effector protein AvrSr50 escapes Sr50 recognition by a substitution in a single surface-exposed residue. New Phytol 234(2): 592–606.

Periyannan S, Moore J, Ayliffe M, Bansal U, Wang X, Huang L, Deal K, Luo M, Kong X, Bariana H, Mago R, McIntosh R, Dodds P, Dvorak J, Lagudah E. 2013. The gene Sr33, an ortholog of barley Mla genes, encodes resistance to wheat stem rust race Ug99. Science 341(6147): 786–788.

Saur IM, Bauer S, Kracher B, Lu X, Franzeskakis L, Muller MC, Sabelleck B, Kummel F, Panstruga R, Maekawa T, Schulze-Lefert P. 2019. Multiple pairs of allelic MLA immune receptor-powdery mildew AVRA effectors argue for a direct recognition mechanism. Elife 8.

Schindelin J, Arganda-Carreras I, Frise E, Kaynig V, Longair M, Pietzsch T, Preibisch S, Rueden C, Saalfeld S, Schmid B, Tinevez JY, White DJ, Hartenstein V, Eliceiri K, Tomancak P, Cardona A. 2012. Fiji: an open-source platform for biological-image analysis. Nat Methods 9(7): 676–682.

Schurmann N, Trabuco LG, Bender C, Russell RB, Grimm D. 2013. Molecular dissection of human Argonaute proteins by DNA shuffling. Nat Struct Mol Biol 20(7): 818–826.

Seeholzer S, Tsuchimatsu T, Jordan T, Bieri S, Pajonk S, Yang WX, Jahoor A, Shimizu KK, Keller B, Schulze-Lefert P. 2010. Diversity at the powdery mildew resistance locus from cultivated barley reveals sites of positive selection. Mol Plant Microbe Interact 23(4): 497–509.

Sievers F, Wilm A, Dineen D, Gibson TJ, Karplus K, Li W, Lopez R, McWilliam H, Remmert M, Soding J, Thompson JD, Higgins DG. 2011. Fast, scalable generation of high-quality protein multiple sequence alignments using Clustal Omega. Mol Syst Biol 7: 539.

Stemmer WP. 1994. DNA shuffling by random fragmentation and reassembly: in vitro recombination for molecular evolution. Proc Natl Acad Sci U S A 91(22): 10747–10751.

Tamborski J, Seong K, Liu F, Staskawicz BJ, Krasileva KV. 2023. Altering specificity and autoactivity of plant immune receptors Sr33 and Sr50 via a rational engineering approach. Mol Plant Microbe Interact 36(7): 434–446.

Zhao H, Arnold FH. 1997. Optimization of DNA shuffling for high fidelity recombination. Nucleic Acids Res 25(6): 1307–1308.

