## Supplementary Methods, table and figures for "Identification of a key gain-of-function residue for effector binding by *in vitro* shuffling of barley *Mla NLR* genes"

### Materials and methods

#### DNA shuffling

To generate recombinant *Mla* genes, DNA shuffling experiments were conducted using a modified protocol (Zhao & Arnold, 1997; Meyer *et al.*, 2014). In brief, the DNA shuffling method includes random fragmentation of the parental genes, recombination (shuffling) of the fragments and reassembly into novel full-length genes, followed by amplification of the recombinant genes. To prepare parental genes for shuffling, the WT *Mla* genes were synthesised (Twist Bioscience) and cloned into the yeast expression vector pGBKT7 (Clontech) using yeast recombinational cloning. The parental genes were then amplified from the pGBKT7 constructs using primer pairs (XZ-163 and XZ-125) (Supplementary Table S1) that bind to the *GAL4 BD* sequence and the ADH1 terminator in the vector, up and downstream of each *Mla* gene respectively. This added 80 bp and 170 bp vector sequences to the 5'- and 3'-ends of the *Mla* genes respectively. The amplified DNA products were cleaned using the Wizard SV Gel and PCR Clean-Up System (Promega). For fragmentation, 4 µg parental DNA was diluted to 50 µl in the digestion buffer containing 50 mM Tris pH 7.5 and 10 mM $MnCl_2$. This digestion mixture was equilibrated at 15°C for 10 min before 0.3 units of DNase I (NEB) was added. The digestion was run at 15°C for 1 min or until the majority of the DNA fragments were between 250 – 1500 bp, followed by incubation at 80°C for 10 min to deactivate DNase I. The digested DNA products were separated using agarose gel electrophoresis and fragments with sizes between 250 – 1500 bp were recovered using the Wizard SV Gel and PCR Clean-Up System (Promega). Fragmentated parental genes were mixed in equal amounts and reassembled in a primer-less PCR using Phusion polymerase (NEB) according to the manufacturer's instructions. The PCR mixture contained 20-30 ng/µl of the parental fragments to which no primers were added. A thermocycling program of 98°C for 1 min; 98°C for 30 s, 42°C for 40 s, and 72°C for 45 s + 1 s per cycle (40 cycles); 72°C for 10 min and hold at 4°C was used. To amplify the assembled genes, 1/100 dilution of the products from the primer-less PCR were added to a new batch of Phusion PCR mixture (NEB) with primer pairs (XZ-155 and XZ-156) (Supplementary Table S1) that bind to the vector sequences flanking the *Mla* genes. These primers allowed amplification via nested PCR, generating re-assembled *Mla* genes with 60 bp end sequences homologous to the pGBKT7 vector. A standard Phusion PCR program of 98°C for 30 s; 98°C for 10 s, 55°C for 30 s, and 72°C for 1 min 45 s (35 cycles); and 72°C for 10 min was performed according to the manufacturer's instructions. The PCR products were analysed using agarose gel electrophoresis and amplified DNAs with sizes of approximate 3,000 bp were recovered from the gel.

#### Yeast library construction and Y2H

The yeast libraries were constructed and the Y2H experiments were performed using a method modified from the Matchmaker® Gold Yeast Two-Hybrid system (Takara Bio). In brief, the WT *Mla* genes and the pools of recombinant *Mla* genes were cloned into the pGBKT7 vector using recombinational cloning in the yeast strain Y187 (Clontech). Successful transformants carrying the WT or recombinant *Mla* genes fused to the *GAL4 BD domain* (BD-MLA) were selected based on yeast colony growth on synthetic dropout (SD) medium [2% (w/v) glucose, 0.139% (w/v) yeast synthetic drop-out medium supplement pH 6, 0.67% (w/v) yeast nitrogen base, 2% (w/v) Bacto Agar] without tryptophan (SD-W). The *AVR* genes were cloned into the yeast expression vector pGADT7-GW using Gateway cloning (Invitrogen) and transformed into the yeast strain Y2HGold (Takara Bio). Transformants carrying each *AD-AVR* construct were selected based on yeast colony growth on SD without leucine (SD-L). For each WT *Mla* or *AVR* construct, five colonies were picked, mixed into freezing medium [1% (w/v) yeast extract, 2% (w/v) peptone, 2% (w/v) glucose and 25% (v/v) glycerol] and stored at -80°C. For the recombinant *Mla* libraries, all colonies were harvested from entire plates and resuspend into freezing medium for storage at -80°C, following colony counting using FIJI (Schindelin *et al.*, 2012).

For mating-based Y2H, yeast strains carrying the *AVR* genes or the WT *Mla* genes were grown in corresponding SD media without agar overnight at 28°C with shaking. Yeast strains carrying the recombinant *Mla* genes were reactivated from storage at -80°C by growing on SD-W at 28°C for 4 days before mating. The *Mla*-carrying yeast strains were mixed with corresponding *AD*-carrying yeast strains to a 1:2 ratio and mated overnight at 28°C on the YPDA plates (1% (w/v) yeast extract, 2% (w/v) peptone, 2% (w/v) glucose, 0.01% (w/v) adenine and 2% (w/v) Bacto Agar). The mated yeast cells were then grown on SD-WL to select and enrich diploid yeast carrying both the BD- and AD- constructs. For interaction assays, the diploid yeast colonies were grown on SD-HWL and yeast growth was evaluated from 4 to 10 days after incubation. For co-transformation-based Y2H, the yeast strain Y2HGold was transformed with BD- and AD- constructs and grown on SD-WL. Interaction assays were conducted as above.

To analyse the sequences of the recombinant *Mla* genes, plasmid DNA was extracted from the yeast colonies using a phenol:chloroform method (Kaiser *et al.*, 1994) and used in colony PCR to identify full-length *Mla* genes. PCR products from amplification of the full-length *Mla* genes were cleaned using the Wizard SV Gel and PCR Clean-Up System (Promega) then sequenced using Sanger sequencing. The sequences were aligned using Clustal Omega (Sievers *et al.*, 2011) and recombination was analysed using Salanto (Schurmann *et al.*, 2013). For full-plasmid sequencing, plasmid DNA was extracted from yeast cultures grown overnight in SD-W without agar at 28°C with shaking. The plasmid DNA extracted from yeast was transformed into *Escherichia coli* and further multiplied, before sequencing with Oxford Nanopore long-read sequencing (Plasmidsaurus).

**Transient gene expression in *N. benthamiana* and cell death assays**

The WT *Mla* or recombinant *Mla* genes were cloned into the pBW517 (Saur *et al.*, 2019) vector using Gateway cloning (Invitrogen). The resulting constructs express MLA fused to a 4xMyc tag (MLA-myc). The *AVR* genes were cloned into the pICH47732 vector using Golden Gate cloning (Engler *et al.*, 2014), generating *AVR-GFP* fusion genes. The constructs were transformed into Agrobacterium strain GV3101_pMP90 by electroporation. For agroinfiltration, the transformed Agrobacterium strains were grown in Luria-Bertani liquid medium containing 15 µg/ml gentamicin and 25 µg/ml carbenicillin at 28°C for 24 h with shaking. The Agrobacterium cultures were harvested, resuspended in infiltration medium (10 mM MES pH 5.6, 10 mM $MgCl_2$ and 150 µM acetosyringone) and mixed to a final $OD_{600}$ = 0.5 for each strain. The mixtures were incubated at room temperature for 2 h before infiltration into 3-4 weeks-old *N. benthamiana* leaves using a syringe. Leaves were imaged and cell death phenotypes were analysed 3-5 days after infiltration.

**Supplementary Table**

**Table S1 Primers used in this study**

| Name | Sequences |
|---|---|
| XZ-163 | GAGAGTAGTAACAAAGGTCAAAGACAGTTG |
| XZ-125 | CTATACCTGAGAAAGCAACCTGACCT |
| XZ-155 | ATGGAGGAGCAGAAGCTGATCTCA |
| XZ-156 | CCCGTTTAGAGGCCCCAAGG |

**Supplementary figures**

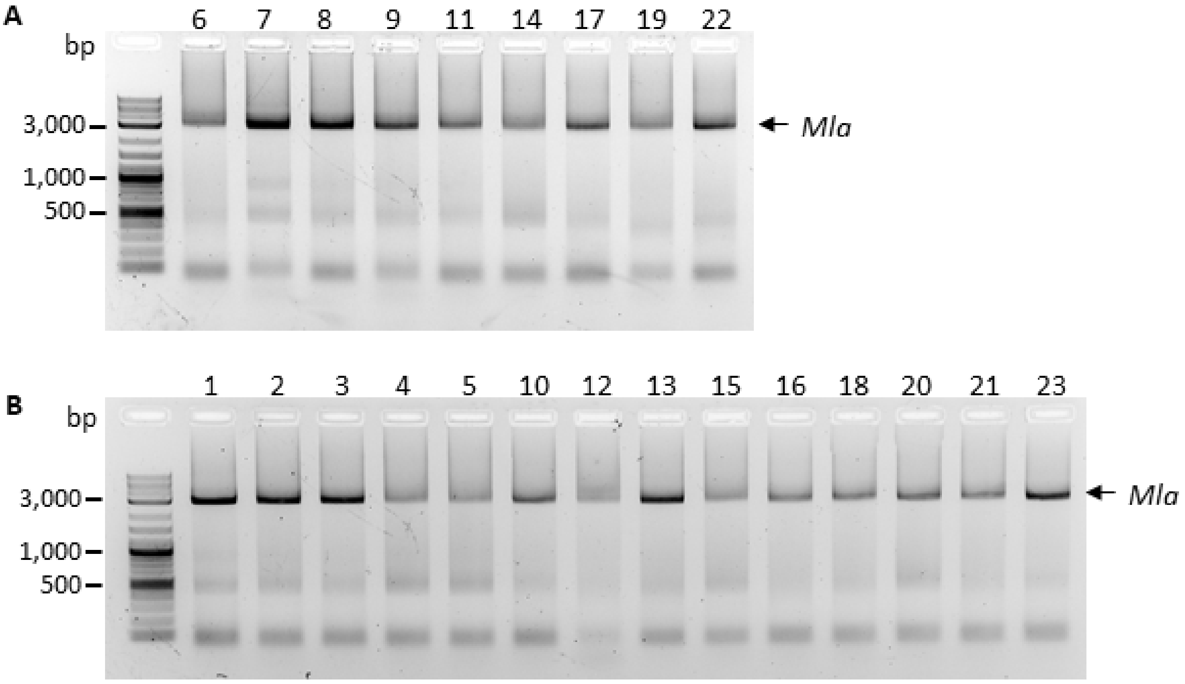


C

| | | Barley | | | | Wheat | | | |
|---|---|---|---|---|---|---|---|---|---|
| | | *Mla7* | *Mla13* | *Mla10* | *Mla1* | *Sr33* | *TmMla1* | *Sr50* | |
| Barley | *Mla7* | | 1 | 2 | 3 | 4 | 5 | 6 | |
| | *Mla13* | | 7 | 8 | 9 | 10 | 11 | 12 | |
| | *Mla10* | | | | 13 | 14 | 15 | 16 | |
| | *Mla1* | | | | | 17 | 18 | 19 | |
| Wheat | *Sr33* | | | | | | 20 | 21 | |
| | *TmMla1* | | | | | | | 22 | |
| | *Sr50* | | | | | | | | |
| | Barley 6 X *Mlas* | *Mla1, Mla7, Mla9, Mla10, Mla13 and Mla22* | | | | | | | 23 |

**Figure S1. DNA shuffling of barley and wheat *Mla* genes.** (A and B) Agarose gel electrophoresis of PCR products amplified from the DNA shuffling experiments. The numbers of the samples are indicated on top of each sample well. The expected size of a full-length *Mla* gene is indicated by an arrow at the right of each gel. (C) Sample numbering for (A) and (B).

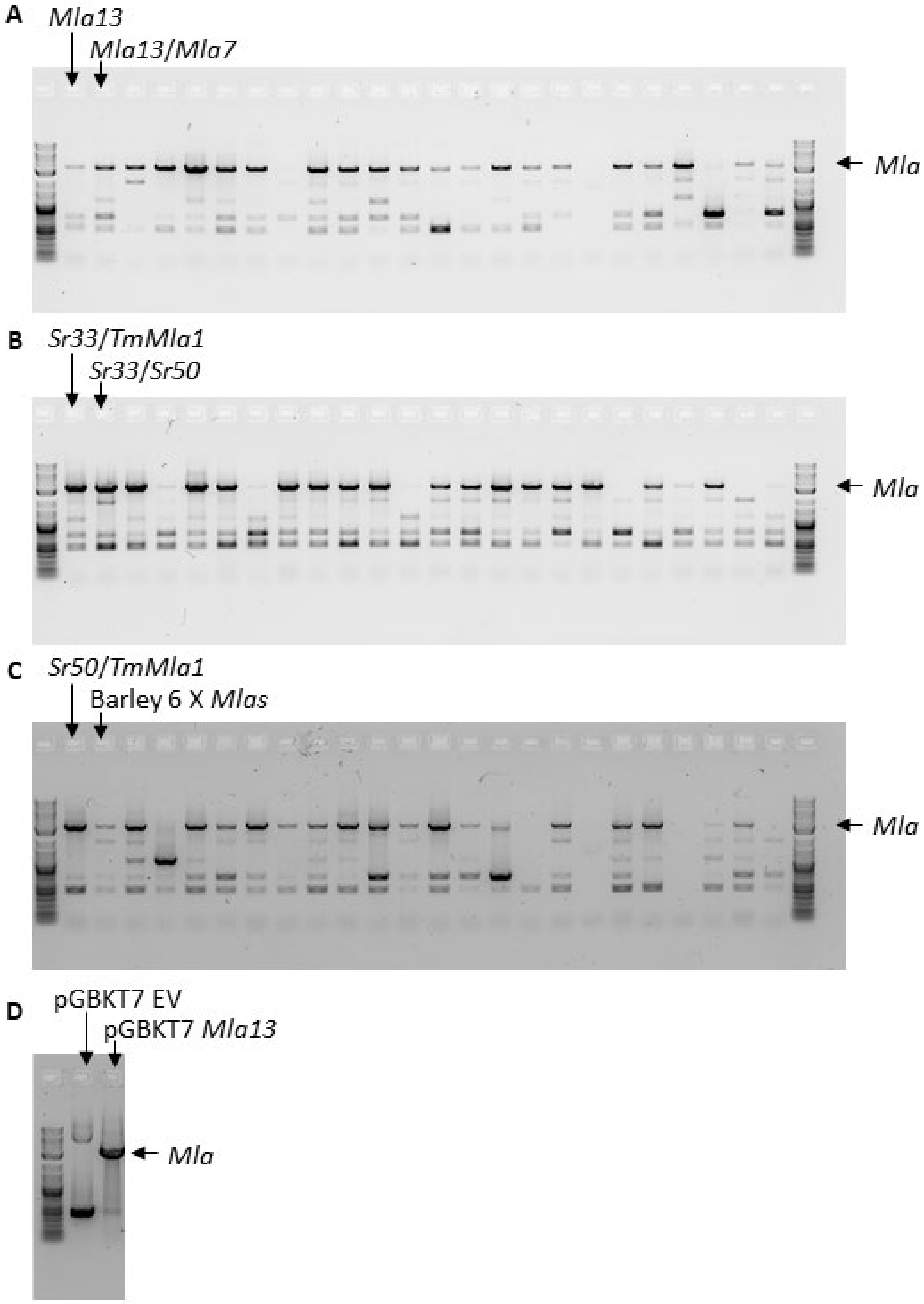


**Figure S2. Colony PCR of yeast libraries for identification of full-length *Mla* genes.** (A-C) Full-length *Mla* genes were identified in colonies picked from six yeast recombinant gene libraries. For each library, 12 colonies were picked randomly and colony PCR products from two libraries were loaded alternately into the sample wells in each gel as indicated on top of the gels. (D) PCR controls using pGBKT7 empty vector (EV) or pGBKT7-MLA13 plasmid DNAs as templates. The expected size of a full-length *Mla* gene is indicated by an arrow at the right of each gel.

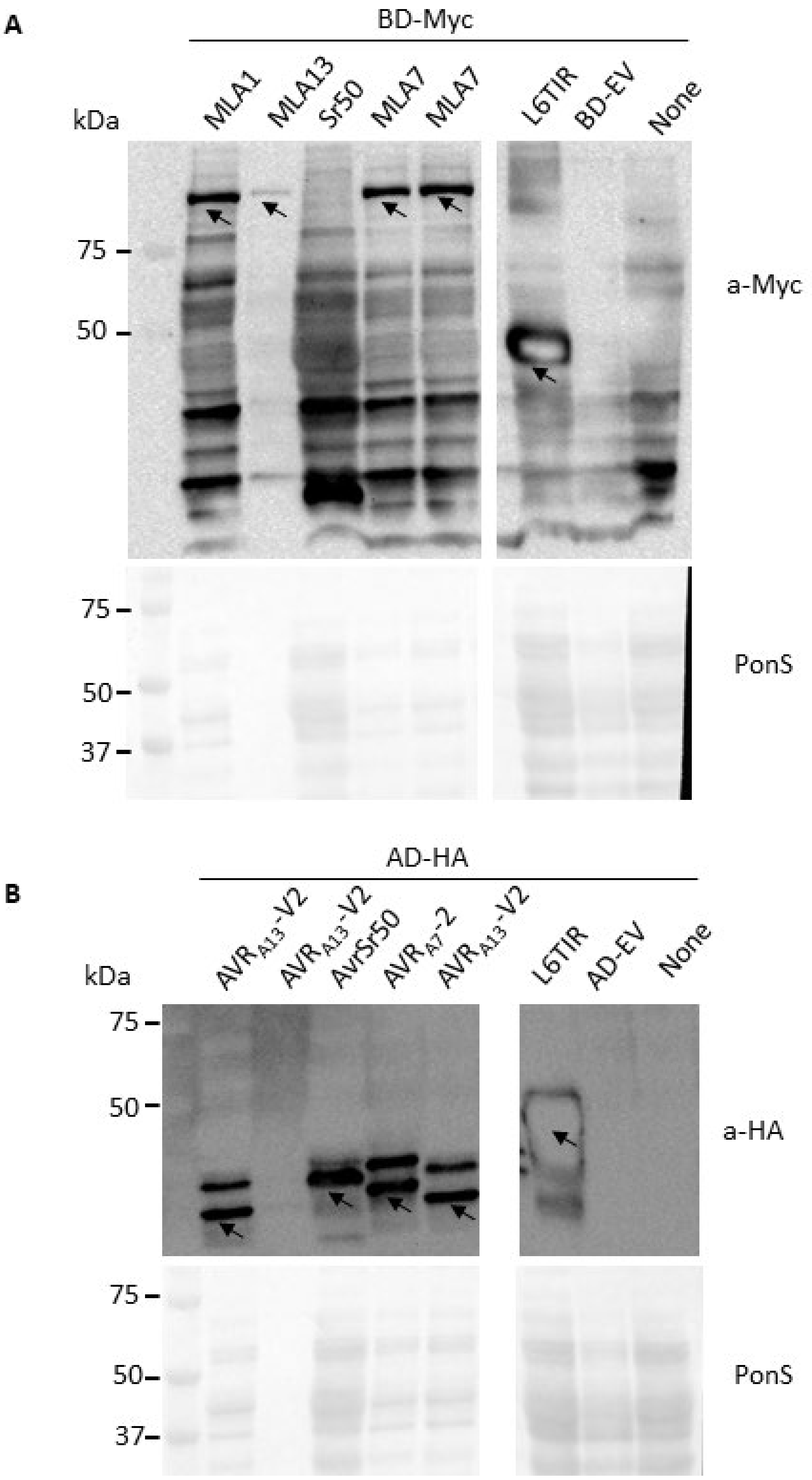


**Figure S3. Western-blot analysis of protein expression in yeast.** Protein extracts from diploid yeast colonies containing both a BD- and an AD- constructs were analysed for the presence of BD-Myc fusion proteins (A) and AD-HA fusion proteins (B). Arrows indicate the presence of the corresponding proteins. Ponceau S (PonS) staining indicates protein loading on gels.

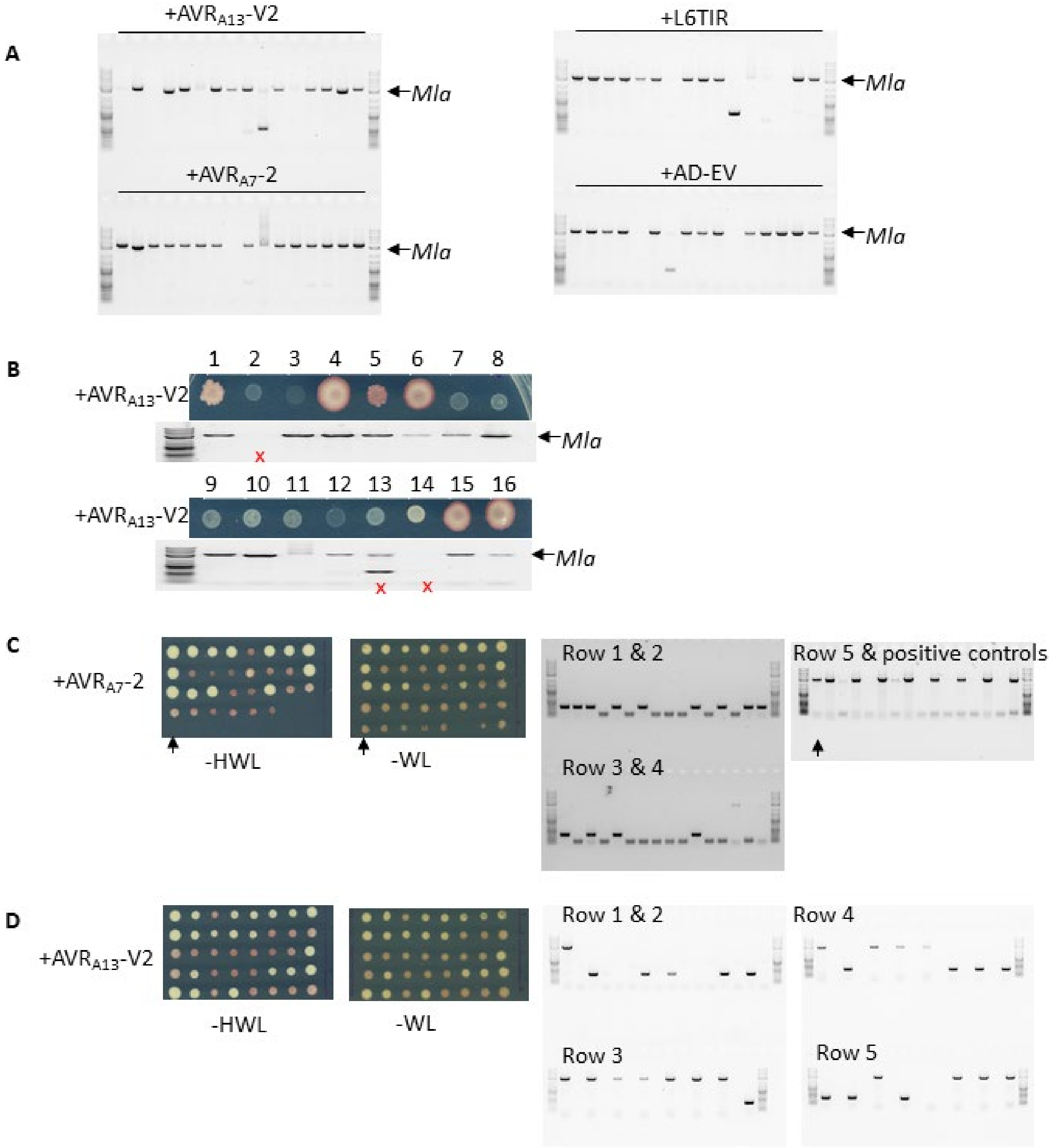


**Figure S4. Full-length *Mla* genes identified from colonies of the *Mla13/7* library mated with various AD-containing constructs.** (A) Colony PCR and agarose gel electrophoresis of the small-scale Y2H assay, in which 16 diploid colonies were randomly picked for each gene combination. (B) Amplification of full-length *Mla* genes from colonies of *Mla13/7* mated with AVR$_{A13}$-V2 for Sanger sequencing. The corresponding colony growth on the interaction plate is shown above the agarose gel. Samples that failed during Sanger sequencing are marked by red crosses. (C and D) Colony PCR and agarose gel electrophoresis of the large-scale Y2H assay, in which 40 colonies were randomly picked from the interaction plates of *Mla13/7* mated with AVR$_{A7}$-2 (C) and AVR$_{A13}$-V2 (D) respectively. The left two panels show replica yeast growth on interaction plates (-HWL) and diploid plates (-WL). The right two panels show agarose gel electrophoresis of the corresponding colony PCR products. The arrow in (C) indicate the colony with a full-length *Mla* but failed to grow in the replica interaction (-HWL) plate.

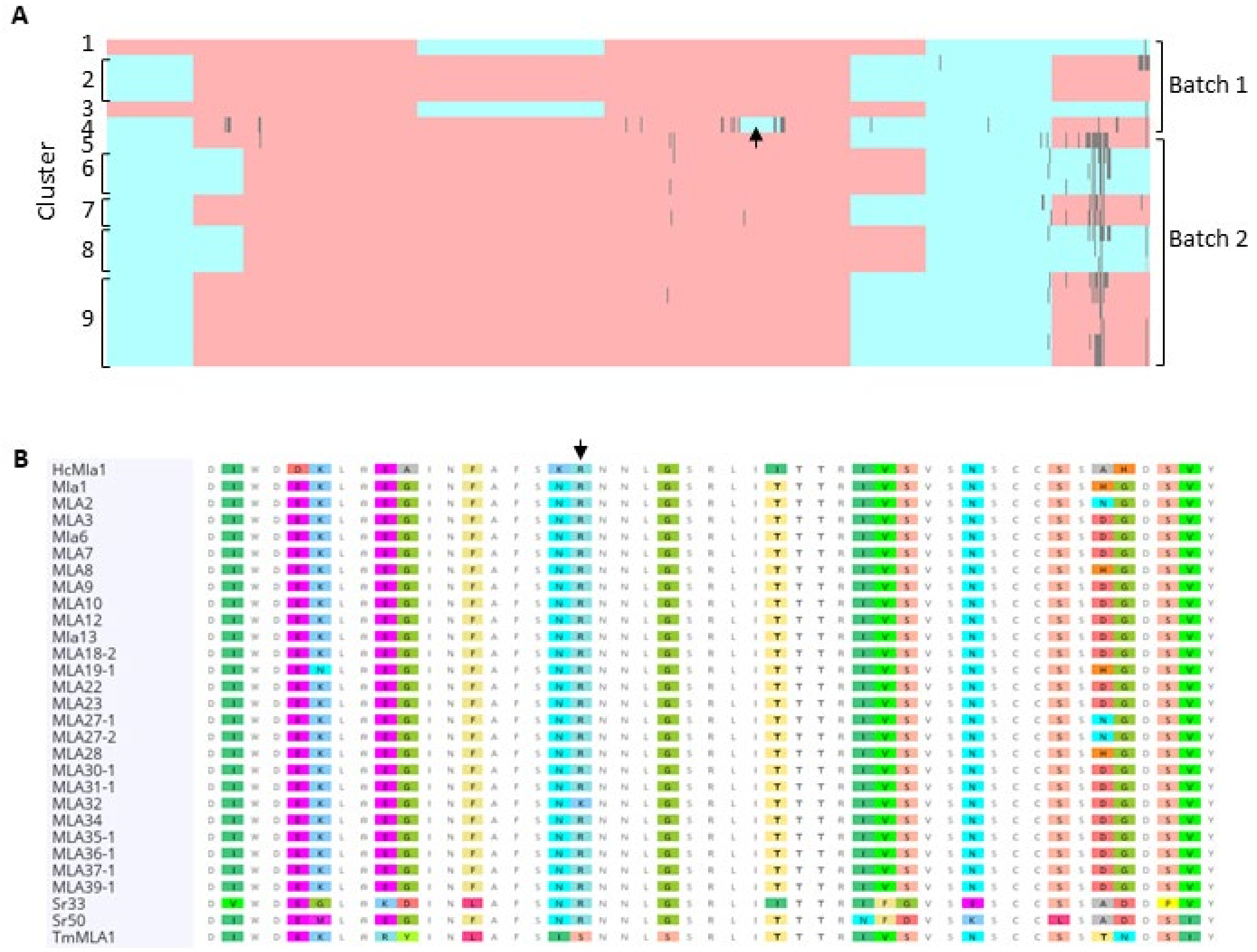


**Figure S5. Sequence alignment and DNA shuffling analysis of recombinant *Mla13/7* genes encoding AVR$_{A13}$-V2 interactors.** (A) A cartoon depiction of sequence alignment and DNA shuffling analysis of 21 recombinant sequences encoding MLA proteins that interacted with AVR$_{A13}$-V2. The gene fragments assigned to *Mla7* and *Mla13* are coloured in red and blue respectively. Mutations that were not assigned to either parent are shown in dark grey. The arrow indicates a synonymous variation. The clusters and the experimental batches of the sequences are showed to the left and right of the alignment respectively. (B) Amino acid sequence alignment of barley and wheat MLA proteins. The arrow indicates the R298 residue that is conserved in most MLA proteins but is a serine residue in TmMLA1.

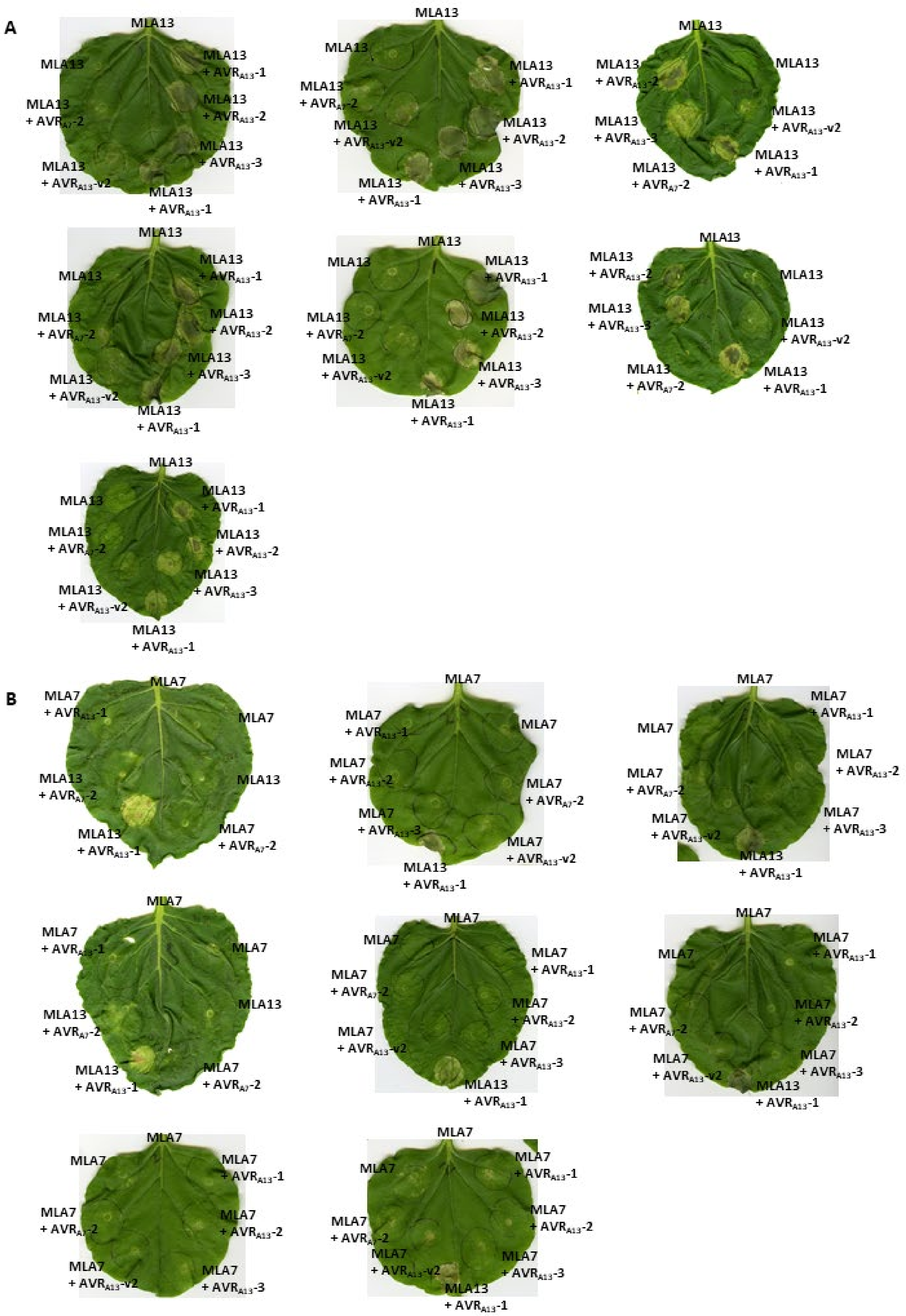


**Figure S6. Cell death assays for MLA13, MLA7 and MLA13/7 variants in *N. benthamiana* leaves.**

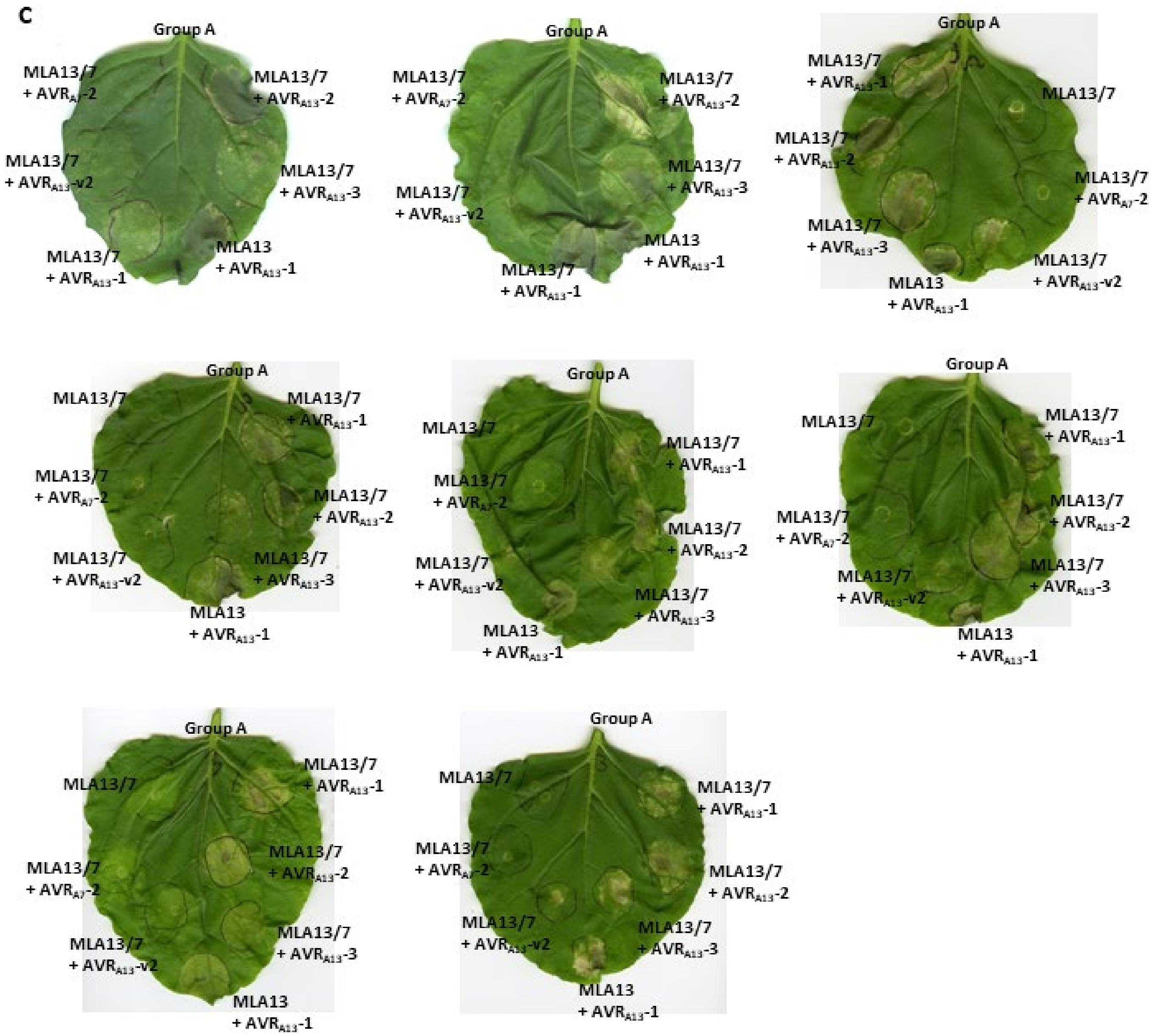


**Figure S6. Cell death assays for MLA13, MLA7 and MLA13/7 variants in *N. benthamiana* leaves.** (continued)

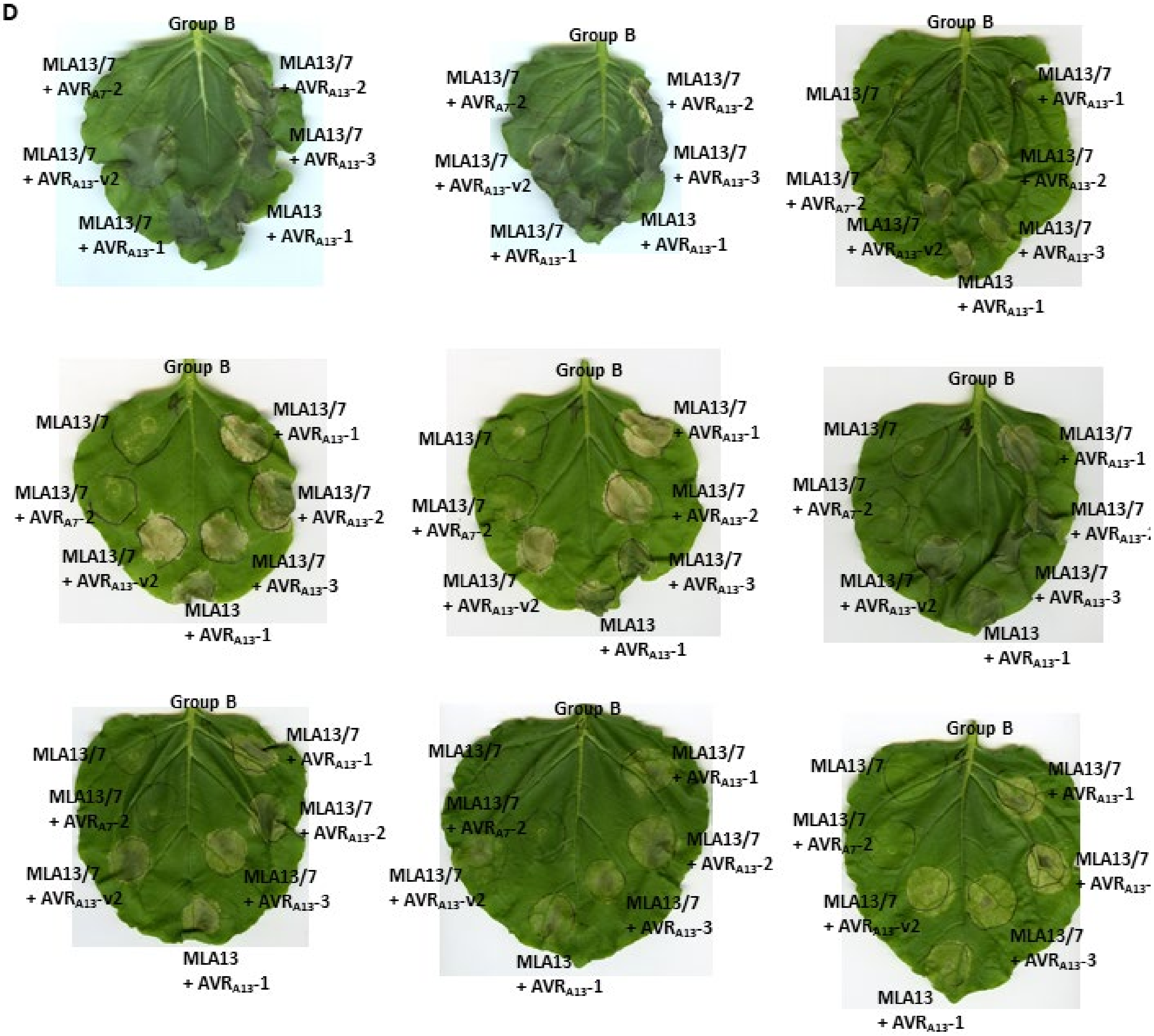


**Figure S6. Cell death assays for MLA13, MLA7 and MLA13/7 variants in *N. benthamiana* leaves.** (continued)

**E**

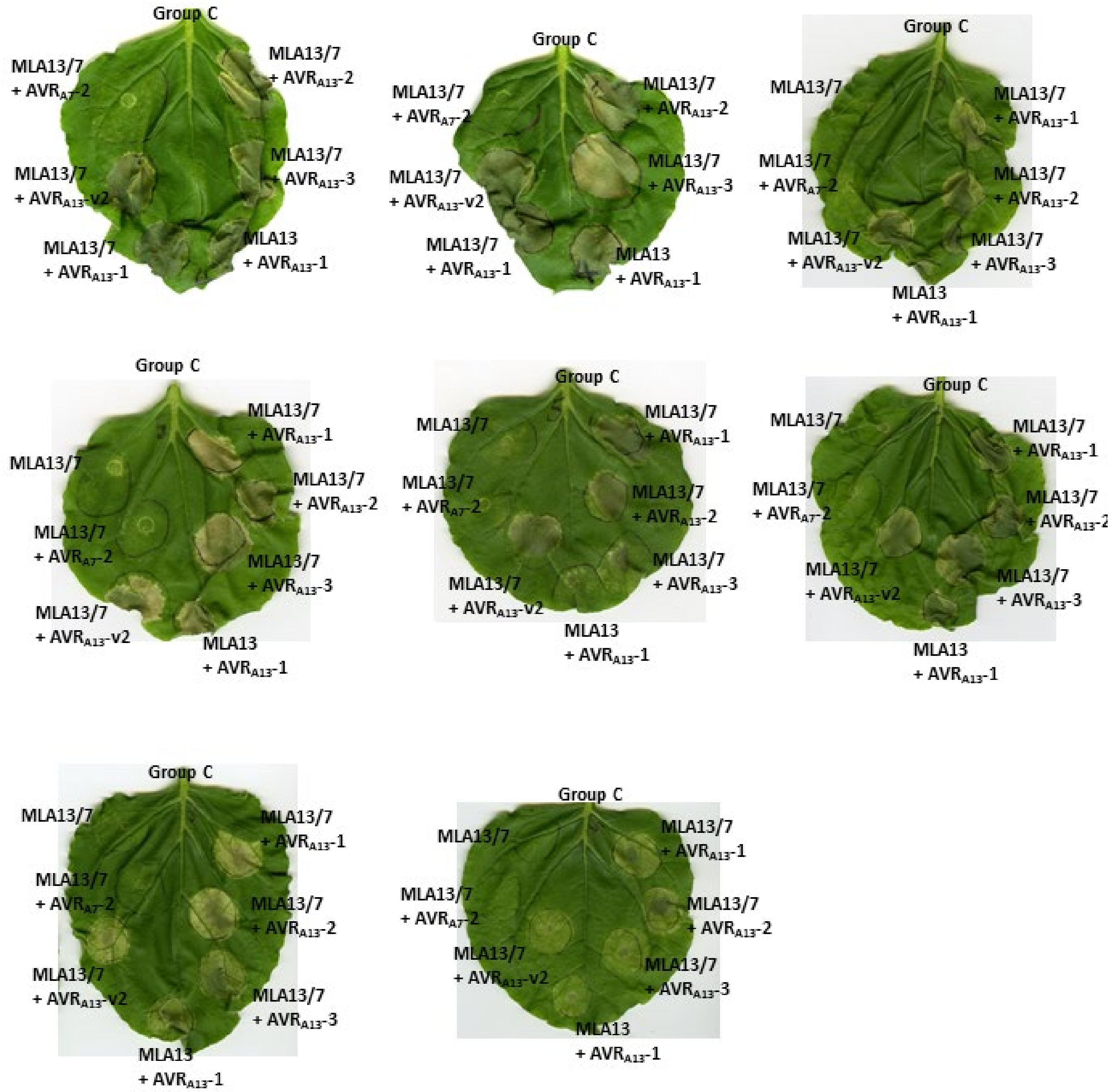


**Figure S6. Cell death assays for MLA13, MLA7 and MLA13/7 variants in *N. benthamiana* leaves.** Agrobacterium strains carrying MLA13 (A), MLA7 (B) or one of the MLA13/7 variants (C-E) were co-infiltrated with Agrobacterium strains carrying one of the AVR$_{A13}$ variant, AVR$_{A7}$-2 or alone into *N. benthamiana* leaves. Images were taken 5 days after infiltration. Yellow to dark spots on the leaves indicate cell death.

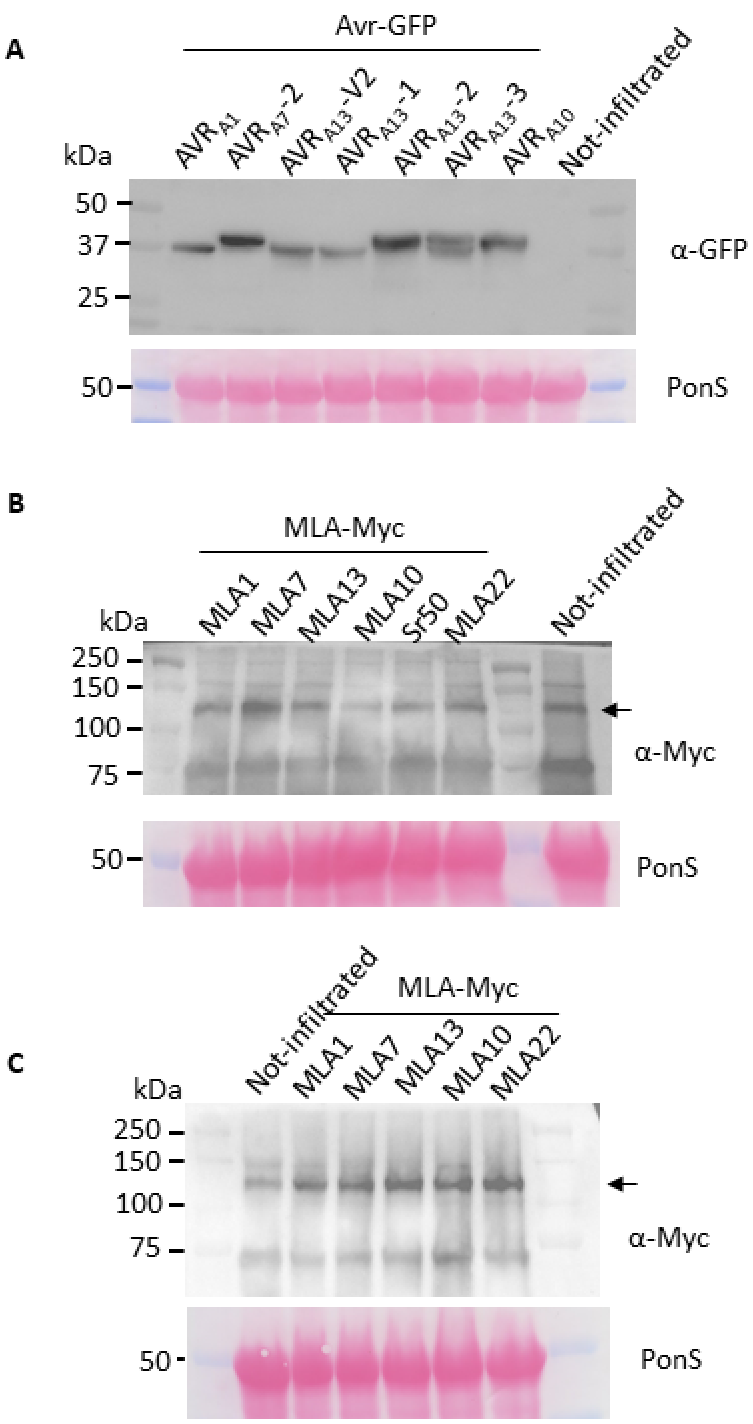


**Figure S7. Protein accumulation in *N. benthamiana*.** Protein accumulation was assessed by western blotting with appropriate antibodies. (A) AVR-GFP fusion proteins. Samples were collected 22 h hours post agroinfiltration. (B and C) MLA-Myc fusion proteins. Samples were collected 22 hours (B) and 72 hours (C) post Agro-infiltration respectively. The arrows indicate the expected size of the MLA-Myc fusion proteins (114 kDa). The 72 h sample was not collected for Sr50 because of the extensive cell death in the infiltrated area at this timepoint.

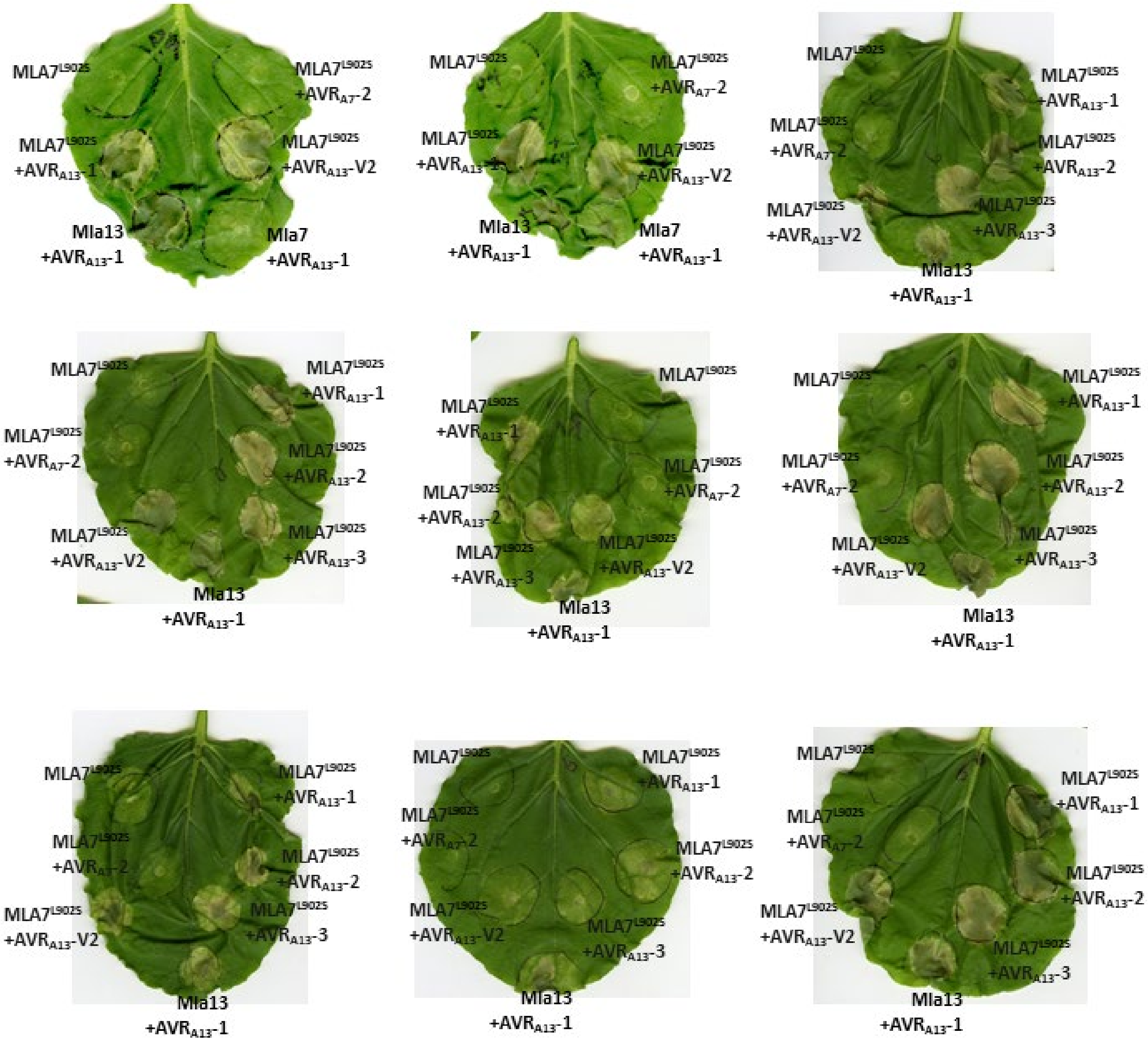


**Figure S8. Cell death studies of the *Mla7$^{L902S}$* mutant in *N. benthamiana* leaves.** Agrobacterium strains carrying *Mla13*, *Mla7* or *Mla7$^{L902S}$* were co-infiltrated with Agrobacterium strains carrying *AVR$_{A13}$-V2*, *AVR$_{A13}$-1*, *AVR$_{A13}$-2* or *AVR$_{A7}$-2* into *N. benthamiana* leaves. Images were taken five days after infiltration. Cell death is indicated by yellow to dark spots on the leaves.
